# A functional module states framework reveals cell states for drug and target prediction

**DOI:** 10.1101/2020.11.24.394932

**Authors:** Guangrong Qin, Theo Knijnenburg, David Gibbs, Russell Moser, Raymond J. Monnat, Christopher Kemp, Ilya Shmulevich

## Abstract

Cells are complex systems in which many functions are performed by different genetically-defined and encoded functional modules. To systematically understand how these modules respond to drug or genetic perturbations, we developed a Functional Module States framework. Using this framework, we 1) defined the drug induced transcriptional state space for breast cancer cell lines using large public gene expression datasets, and revealed that the transcriptional states are associated with drug concentration and drug targets; 2) identified potential targetable vulnerabilities through integrative analysis of transcriptional states after drug treatment and gene knockdown associated cancer dependency; and 3) used functional module states to predict transcriptional state-dependent drug sensitivity and built prediction models using the functional module states for drug response. This approach demonstrates a similar prediction performance as do approaches using high dimensional gene expression values, with the added advantage of more clearly revealing biologically relevant transcriptional states and key regulators.

## Introduction

Cells are complex systems that have been investigated at multiple levels, including genomic, epigenomic, transcriptomic, proteomic and metabolomic. The varying concentrations and abundances of molecular species reflect diverse processes that regulate cell function. On the genomic level, tumors are often stratified into different subtypes according to the mutation status of certain genes, which are known to be predictive of clinical outcomes (Papaemmanuil et al., 2016; Schmitz et al., 2018). The transcriptome, however, can be affected by genetic and epigenomic alterations and is a more direct lens to investigate cell behavior and provide clues to understand tumor heterogeneity or drug response.

A range of methods have been developed to measure the transcriptome at different levels of resolution, such as gene expression microarrays, bulk RNA sequencing (RNASeq), the L1000 platform (Subramanian et al., 2017) and single cell RNASeq (Zheng et al., 2017). These high-throughput techniques have been used to capture the transcriptomes from thousands of primary tumor samples and cell lines. For example, the TCGA (Knijnenburg et al., 2018; Thorsson et al., 2018) project measured over 10,000 tumor samples for 33 tumor types using RNASeq, the Connectivity Map (Subramanian et al., 2017) project provided over one million transcriptomic profiles of different cell line samples after treatment of drugs or knockdown of genes using the L1000 platform, and the GDSC (Iorio et al., 2016) and CCLE (Barretina et al., 2012; Ghandi et al., 2019) projects measured transcriptional profiles of multiple cell lines before drug treatment using microarrays or RNASeq. The growing archive of transcriptomic data provides a rich source of information for defining transcriptional states and understanding the functionality of cells that are perturbed by genetic alterations or drug treatment.

A number of methods have been developed to determine cellular states, especially in the area of single cell studies, such as Monocle (Trapnell et al., 2014), scEpath (Jin et al., 2018), SLICE (Guo et al., 2017), SCENT (Teschendorff and Enver, 2017), and OncoGPS (Kim et al., 2017) which works with bulk data. The main differences among these methods are reflected in the underlying assumptions. Depending on the assumption, different factors, which are based on the biological questions and contexts, are chosen to interpret the cellular transcriptomic state. Commonly used factors include principal components (Tsuyuzaki et al., 2020), entropy (Jin et al., 2018) and various oncogenic transcriptional signatures (Kim et al., 2017). The major shortcomings of current methods to define transcriptional states include: 1) dimensionality reduction without using existing biological knowledge, which can hamper interpretation of cell states; 2) not defining cellular functionalities; 3) using only one factor to represent the transcriptomic states. To overcome these shortcomings, we proposed a functional module states framework to define cell states by gene set or pathway derived factors, which are numeric values that estimate the overall activity of the pathway or gene set.

Curated pathways, such as KEGG (Kanehisa and Goto, 2000; Ogata et al., 1999), cover a large number of genes and a diversity of biological processes, including DNA replication, transcription, energy metabolism, signaling, and others. The expression of genes in these pathways may provide one way to estimate the activity of such functional modules. We proposed the states of cells can be defined by the overall activity profile of the functional modules, which we define as functional module factors (FM-factors). The vector of these so-called FM-factors is then used to represent the functional module states (FM-States). In the remainder of this work, we demonstrate the utility of the newly established FM-States framework by addressing the following questions, 1) are the transcriptomic states of a cancer cell line (MCF7) after drug treatment associated with drug concentration or the drug targets?; 2) can we predict targetable vulnerabilities using the transcriptional cell states?; and 3) can we use the functional states of cancer cell lines (multiple breast cancer cell lines) prior to drug treatment to predict the drug response? We have chosen to focus on breast cancer, but this approach is general and can be readily applied to other tumor types.

## Results

### Overview of the FM-States framework

The goal of the functional module (FM) states framework is to define biologically interpretable factors from high dimensional gene expression data. The resulting FM-factor matrix can be used for further clustering, annotation, detecting regulators for cell states and supervised machine learning, such as classification. The overview of the FM-states framework is shown in Fig 1. We considered each of the pathways in KEGG (Kanehisa and Goto, 2000; Ogata et al., 1999) as a functional module. These functional modules cover diverse cellular activities under the categories of metabolism, genetic information processing, environmental information processing and cellular processes. For each functional module, we defined four types of factors, namely *ssGSEA_score*, *up_strength*, *down_strength*, and *TF_strength*. ssGSEA_score is used to represent the ranking of the expression level for different functional modules in a given sample (Hanzelmann et al., 2013). The *up_strength* and *down_strength* factors estimate the ratio of genes showing high expression (95th percentile or z-score >1.6) or low expression (5th percentile or z-score < −1.6) in one functional module for one specific sample, as compared to all other samples. *TF_strength* estimates the weighted average expression level of transcription factors that are predicted to regulate genes in a given functional module (see methods). After defining the FM-factor matrix, consensus clustering (Monti et al., 2003; Wilkerson and Hayes, 2010) can be used to determine the number of clusters (states), given the input samples. This framework provides a clear biological interpretation for each state by annotating states with different factors and enriched transcription factors (see methods). The functional module factors can also serve as features for machine learning methods, which can identify the key features (i.e., FM-factors) that distinguish different phenotypes or responses to perturbations, such as gene knockdowns or drugs, as we demonstrate.

**Fig 1.**
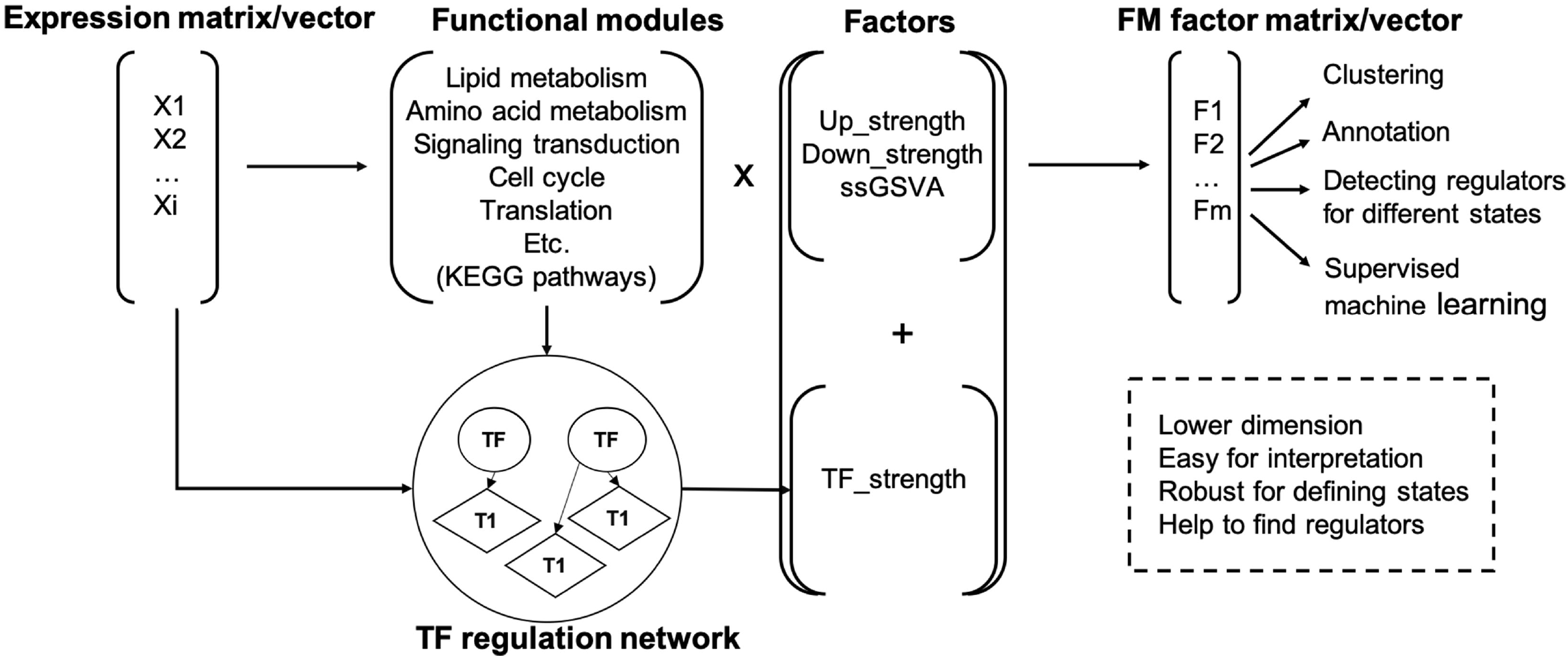
Overview of FM-States framework.

### Functional transcriptional states of breast cancer cell lines reflect drug response

We applied the FM-States method to the high-throughput transcriptional profiles of MCF7 cell lines after drug treatment available from Cmap (Subramanian et al., 2017), following the pipeline shown in supplemental Fig S1. MCF7 cells with available drug sensitivity data (EC50 values) from the Cancer Therapeutics Response Portal (CTRP2) were selected (Aksoy et al., 2017; Basu et al., 2013; Rees et al., 2016; Seashore-Ludlow et al., 2015), resulting in a dataset of 1287 gene expression profiles that represent the transcriptome of MCF7 cell line treated with 190 drugs or compounds in different concentrations (drug-treated samples). A reference gene expression dataset of 1400 MCF7 samples treated with DMSO or H_2_O from CMap (Subramanian et al., 2017) was also used (drug-free samples).

Twenty-three functional modules (supplemental table 1) and the four categories of factors including *ssGSEA_score*, *up_strength*, *down_strength*, and *TF_strength* were selected to define the functional states of MCF7 cells. FM-factors associated with drug treatment were selected by comparing the drug treated samples and drug free samples(Mann-Whitney rank test followed by Benjamini-Hochberg adjustment, FDR <1e-6 and effect size >0.2 (up-regulated) or effect size <-0.2 (down-regulated)). The FM-factors including ssGSEA scores of cell cycle, replication and repair, nucleotide metabolism, transcription and translation were significantly down-regulated after drug treatment, while membrane transport, signal transduction, and lipid metabolism were upregulated after drug treatment (Supplemental Fig S2). FM-factors for the drug treated samples were normalized using reference samples without drug treatment. Then, the drug treatment associated FM-factors were selected for further analysis with consensus clustering to define potential states after drug treatment.

Five transcriptional states were detected for MCF7 cells after drug treatment (Fig 2A, Supplemental Fig S3). Compared to drug-free samples, state 1 (S1) shows up-regulation of the modules involved in replication and repair, transcription, translation and cell cycle, and down regulation of modules involved in transport and catabolism, signaling molecules and interaction, carbohydrate metabolism, and membrane transport (**Active cell cycle state,** Fig 2A & 2B). States 2 and 3 (S2 & S3) show no significant difference compared to drug-free samples (**Basal state**), though S3 shows a somewhat lower expression of cell cycle related modules (Fig 2A & 2B). Both state 4 (S4) and state 5 (S5) featured up-regulation of transport and catabolism, carbohydrate metabolism, signal transduction, amino acid metabolism, and apoptosis, and down-regulation of translation, transcription, replication and repair, nucleotide metabolism, cell cycle and apoptosis. Specifically, S4 features significant down-regulation of transcription factors that regulate replication and repair, cell cycle and cellular senescence (**High apoptosis**/**Low cell cycle/Cell cycle TF suppressed,** Fig 2C). S5 features up-regulation in the transcription factors that regulate various functional modules including transport and catabolism, lipid metabolism, replication and repair, cell cycle, and cell motility (**High apoptosis/Low cell cycle/Cell cycle TF activated**). In this way, the FM-States framework provides direct annotation for different states by measuring the FM-factors among different states.

**Fig 2.**
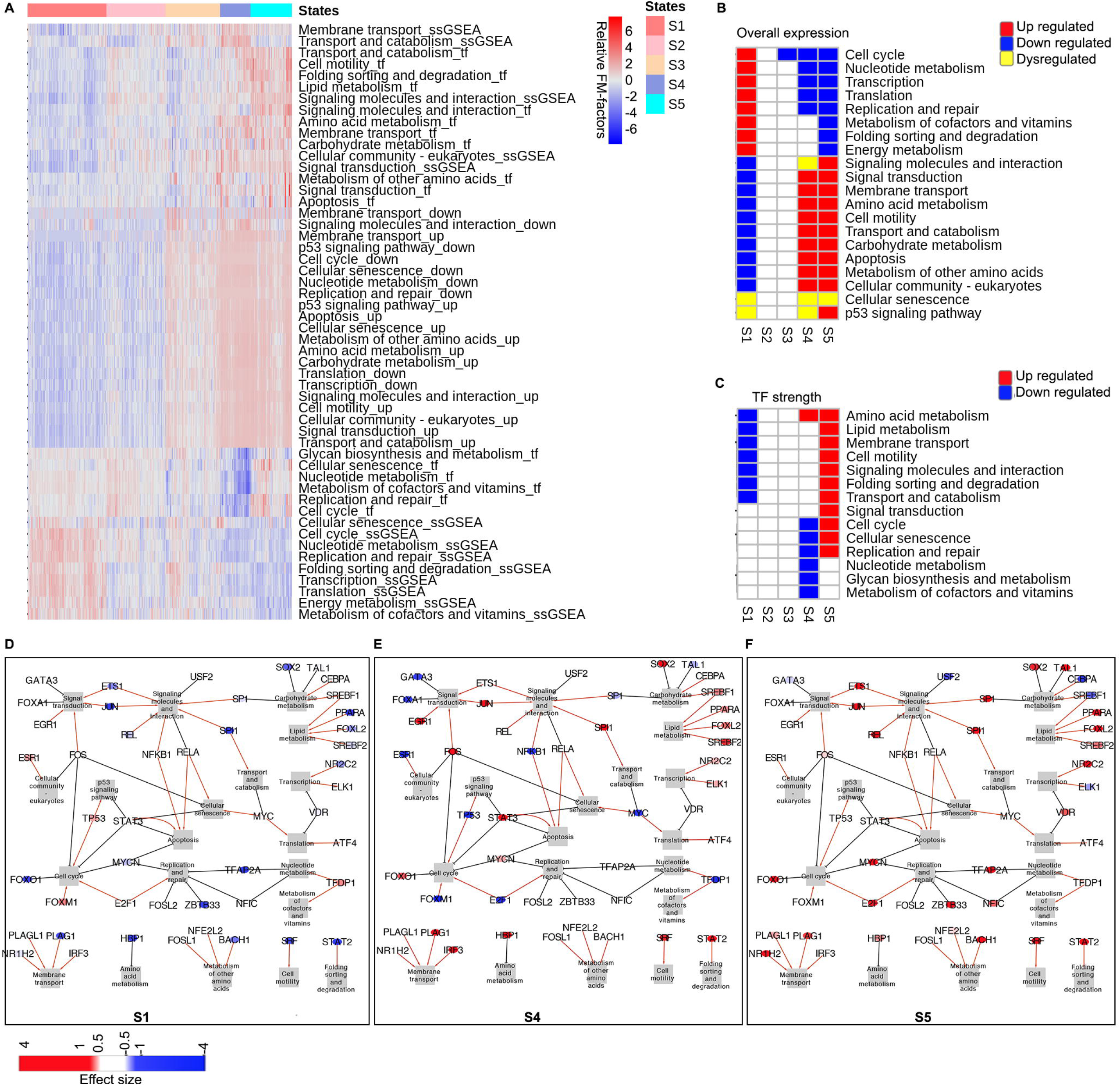
Five states were revealed for MCF7 breast cancer cell line transcriptome after drug treatment using the FM-Factors and transcriptional factors. A. Heatmap of relative FM-factors for MCF7 breast cancer cell lines after treatment of different drugs in different concentrations normalized by the reference MCF cell line without drug treatment. Five states are defined by consensus clustering methods. X-axis shows the 1287 drug treated MCF7 cells; y-axis shows the relative FM-factors scaled by row. B. Functional modules with FM-factors (ssGSEA, up-strength and down-strength) showing significant differences between one state to at least three other states using Wilcoxon rank-sum test in the scipy.stats library (*P* <0.01 is considered as threshold for significance here), and with size effect greater than 1 or smaller than −1(see methods). Functional modules with effect size between one state and all others greater than a threshold (Effect size >1) in the category of ssGESA, up-strength, or smaller than −1 in the down-strength are considered up-regulated; effect size smaller than −1 in ssGESA, up-strength, or greater than 1 in down-strength are considered as down-regulated. The label of dysregulated represents a module with alteration on both sides. C. Differential TF strength among different states using the same threshold as Fig 2B. D-F. Transcriptional regulatory network which may drive the transcriptional states, D(S1), E(S4), F(S5). Transcriptional factors which show enrichment of their target genes in the functional module that show significant difference in the FM-factors among different states, and the transcriptional factors themselves show differential expression between each state to all the other states are shown in D, E and F. The red color represents highly expressed (Effect size >1), and the blue color represents low expressed (Effect size <-1). Arrow means activation (enrichment of positive correlation target genes for the TF in the functional module), and the solid hammer line means suppression (enrichment of negative correlation target genes for the TF in the functional module).

The FM-States framework also provides annotations for the FM-States by detecting the transcription factors that may regulate different states. We annotated the activation or inhibitory effects of one transcription factor on a pathway by the enrichment of its target genes with positive or negative correlation in the pathway. Fig 2D-F shows the transcriptional factors with (1) significant differential expression among different states (one-way ANOVA test, and Bonferroni multiple test correction) and (2) their target genes enriched in the differentially regulated functional modules. For example, FOXO1 shows inhibitory effects to cell cycle, and it shows lower expression in S1 (Fig 2D) and higher expression in S4 and S5 (Fig 2E & 2F). This is consistent with the observation that S4 and S5 are low-cell cycle states. It is also supported by a previous study which showed FOXO1 was associated with cell cycle inhibition (Schmidt et al., 2002). The basal-like state (S2 & S3) didn’t show significant differences in transcription factor expression. Expression of JUN is reduced in S1 and increased in S4 and S5 (Supplemental Fig S4).

Previous time-course studies of MCF7 cells after drug treatment showed the expression of JUN gene elevated after 36h after treatment of chemotherapeutic agents such as doxorubicin and 5-fluorouracil (Troester et al., 2004). SOX2 also shows up-regulation in S4 and S5 (Supplemental Fig S4). Upregulation of SOX2 has been reported to promote the cancer stem cell-like phenotype associated with resistance upon anti-cancer drug treatment (Huser et al., 2018). To further annotate the effects of each transcription factor to each module, we performed enrichment analysis (Fisher’s exact test) to estimate whether there is significant enrichment of the positive correlated target genes or the negatively correlated genes. If the positively correlated target genes for one transcription factor were enriched in one module, we define this transcription factor has an activation effect to this module. Otherwise, if the negatively correlated target genes for one transcription factor are enriched in one module, we define this transcription factor has an inhibitory effect to this module. For example, the enrichment analysis of the MYC target genes which are positively correlated with the expression of MYC are enriched in Cellular Senescence pathway, suggesting MYC plays an activation effect to the module of cellular senescence (Fig 2E, shown in red line). Specifically, MYC shows reduced expression in S4 (Fig 2E, shown in blue color).

Previous studies have suggested that the suppression of MYC induces cellular senescence in tumors (Wu et al., 2007). This result shows the down-regulation of MYC may drive S4 as a cellular senescence state. Similarly, FOXM1 and E2F1, which show activation effects for cell cycle, and replication and repair functional modules are lowly expressed in S4, which may drive the formation of the low cell cycle state of S4. On the contrary, for S5, FOXO1, TFAP2A, ZBTB33 and NFIC, which show inhibitory effects on cell cycle, replication and repair are highly expressed. Previous studies have shown overexpression of the TFAP2A encoded transcription factor AP-2α triggered apoptosis (Muller et al., 2004). The difference of the regulators for the two apoptosis associated states S4 and S5 may suggest different biological mechanisms induced by drugs.

### Functional transcriptional states reveal dose-dependent responses and mechanisms of action

We next set out to determine whether the identified transcriptional states are associated with pharmacologic effects. As the MCF7 cells were treated with different drugs over a range of concentrations, we first asked whether the transcriptional states were associated with drug concentration. By mapping the drug concentration for a specific drug to the drug response curve measured in the CTRP2 project (Rees et al., 2016), we categorized the MCF7 transcriptome profile following treatment of each drug to the high-dosage-drug treated group if the concentration is greater than the EC50 concentration, or the low-dosage treated group if less than the EC50. Results show that the five states induced by drugs are associated with drug concentrations (Chi-Squared test, *P*-value <0.001). Specifically, states S1 and S2 are more enriched in cells treated with low drug concentration, with drug concentration in S3 being higher, and states S4 and S5 being enriched in cells treated with high drug concentration (Fig 3A & 3B). Fig 3A shows the example of doxorubicin. The results show that different drug concentrations induce different transcriptional states.

**Fig 3.**
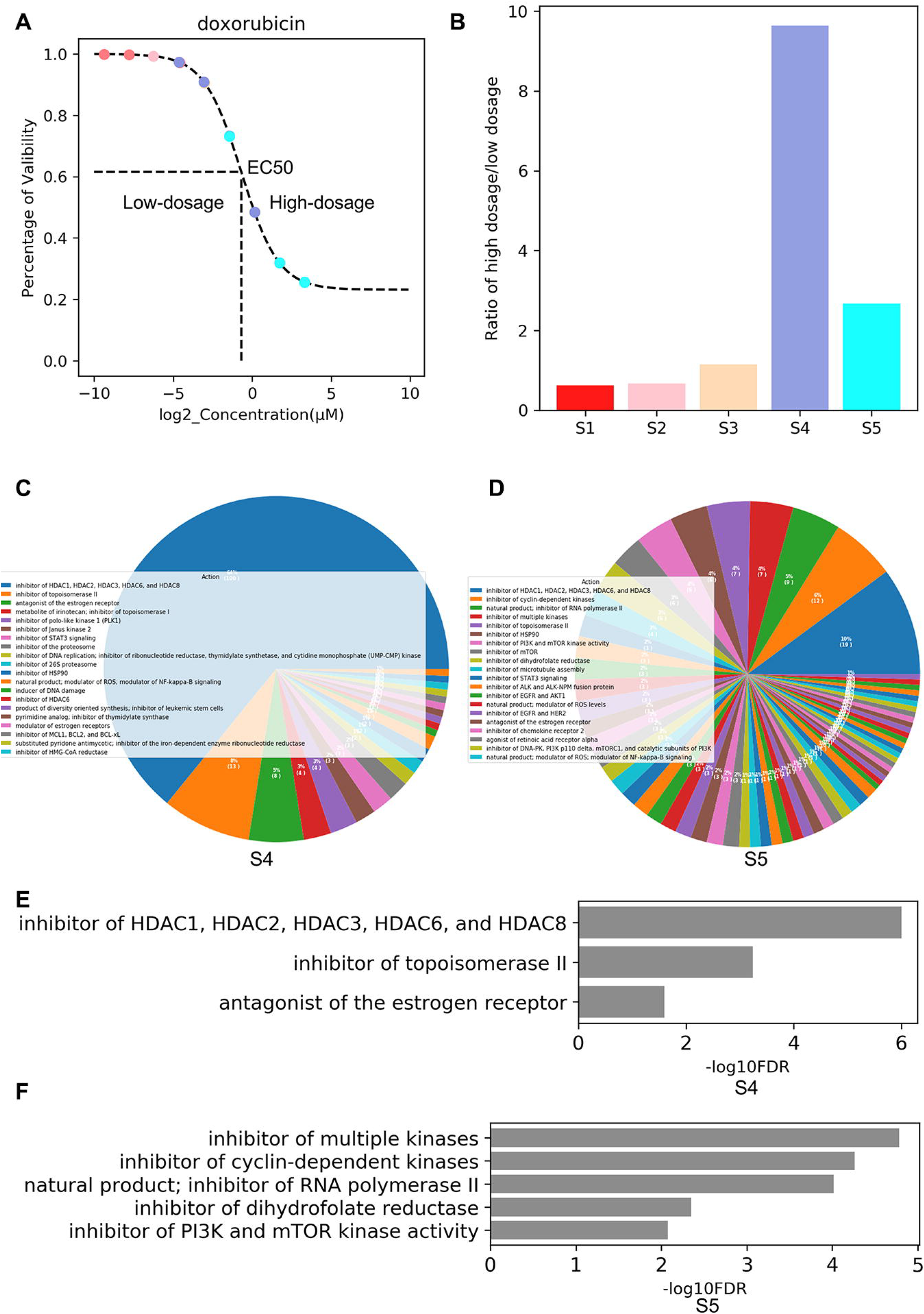
Annotation of MCF7 states after drug treatment using drug dosage and drug targets. A, Mapping of MCF7 cell line profile treated with Doxorubicin in different concentrations to the drug response curve of Doxorubicin for MCF7 as measured in the CTRP2 project, samples treated with concentration greater than greater than EC50 are considered as high-dosage samples, and samples treated with concentration smaller than EC50 are considered as low dosage samples. Samples with different states are colored the same as Fig 3B. B. Histogram of the ratio of high dosages samples over low dosage samples for all drugs compared to the EC50 value for each drug. C-D. Percentage of samples treated with drugs with different targets or actions. C(S4), D(S5). E-F. Enrichment of drug targets for each state using the one-sided Fisher exact test. Results are shown which show *P*-value smaller than 0.05.

We next asked whether drug induced transcriptional states are associated with drug targets or drug action. MCF7 cells have been classified as derived from a Luminal A subtype of breast cancer (Subik et al., 2010), featuring hormone-receptor (estrogen and progesterone-receptor) positive, HER2 negative, and carrying a *PI3KCA* p.E545K mutation (from CCLE and GDSC datasets) (Ghandi et al., 2019; Iorio et al., 2016). Grouping the drugs according to their targets or activity (Iorio et al., 2016), we analyzed the FM states of MCF7 cells treated with drugs (Fig 3C & 3D). The results show that state S4 is enriched in cells treated with estrogen receptor antagonists, topoisomerase II (*TOP2*) inhibitors and *HDAC* inhibitors (Fisher’s exact test, Benjamini-Hochberg multiple comparison(BH adjust), *FDR* <0.05, Fig 3C & 3E). The enrichment of estrogen receptor antagonists, and the down expression of *ESR1* in S4 suggests a drug responsive state. *HDAC* inhibitors play important roles in epigenetic regulation, inducing death and apoptosis and cell cycle arrest in cancer cells (Kim and Bae, 2011). The enrichment of the known approved anti-breast cancer drugs is also consistent with the observation that S4 features high apoptosis and low cell cycle. State S5 is enriched in cells treated with multiple kinase inhibitors, cyclin-dependent kinases inhibitors, inhibitors of RNA polymerase II, inhibitors of dihydrofolate reductase, and inhibitors of PI3K and mTOR kinase activity (Fisher’s exact test, BH adjust, *FDR* < 0.05, Fig 3D & 3F). Since MCF7 cells contain the *PI3KCA* p.E545K mutation, the enrichment of PI3K inhibitors and the alteration in signaling and transduction in S5, suggests S5 is also a drug responsive state. These results suggest that we are able to detect state changes associated with the application of drugs or chemicals with different concentrations and different actions. The FM States annotation of the MCF7 transcriptomic states after drug treatment allows us to achieve more interpretability of the drug induced cell states.

### Functional module states predict therapeutic vulnerabilities

With the classification of transcriptomic states of MCF7 cells after drug treatment, we then asked whether we could use the transcriptomic states to predict therapeutic vulnerability. We made the assumption that the knockdown or knockout of genes that drive drug-responsive transcriptomic states would suggest that these genes are potential therapeutic targets. Using the CMap dataset (Subramanian et al., 2017), which includes transcriptome data following shRNA knockdown, we generated FM-factors and classified each sample into the pre-defined drug-induced states (S1-S5). K-Nearest Neighbor (KNN) classifier was used for the classification. We first assigned each transcriptome profile into one state using the KNN method (Supplemental Fig S5). As each sample represents the transcriptome after gene knockdown using one shRNA seed, we next performed enrichment analysis (Fisher’s exact test, p <0.05) of shRNA seeds for each gene in different states and assigned genes into different states (Fig 4A). To further validate the potential of target dependency, we integrated cancer dependency data from the DepMap project (Dempster et al., 2019; DepMap, 2019; Meyers et al., 2017), which measured the gene knockout effect using CRISPR. Our results show that the MCF7 cell line shows greater sensitivity to depletion of genes that are associated with S3, S4 and S5-like states (Kolmogorov-Smirnov test, *P*-value <0.05) (Fig 4B & 4C). We then used the knockout effect and the proportion of seeds that drive the states as a threshold for selecting the potential gene targets for inducing each state (Supplemental Table 2).

**Fig 4.**
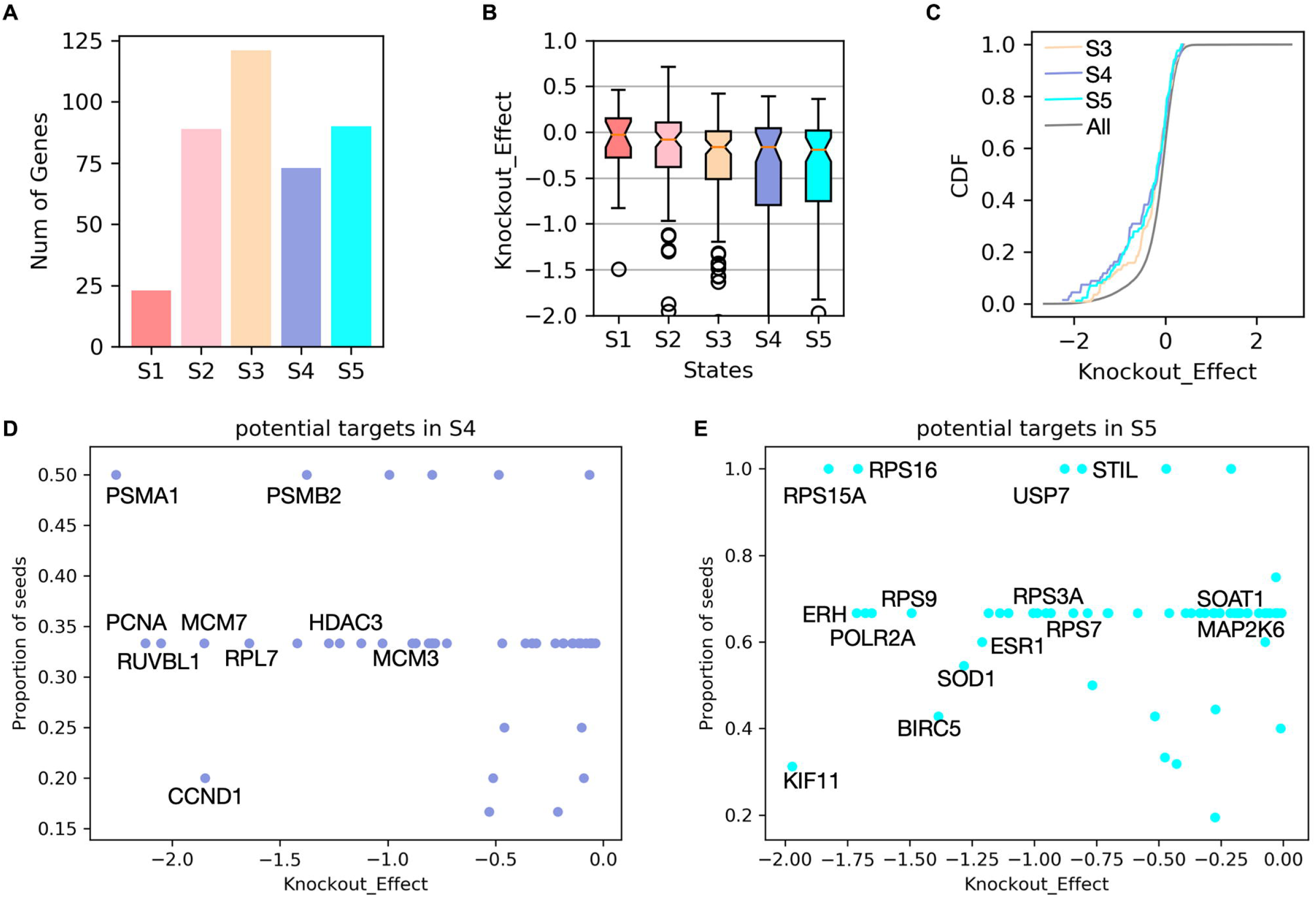
Predicting potential targets for Breast cancer (MCF7 cell line-like). A. Number of genes which are predicted as the knockdown of these genes are associated with each state. B. Boxplot of the gene knockout (KO) effects of state associated genes(Data from Depmap 2019 Q3, Achilles_gene_effect. Median nonessential KO effect is scaled to 0 and the median essential KO effect is −1). C. Cumulative density curve for gene knockout effects in different states, the curve for ‘ALL’ represents the cumulative density curve for the gene knockout effects for all the genes measured in the Depmap project. D, Predicted targets associated with S4 like states, x-axis shows the knockout effect as measured in Depmap data, y-axis shows the proportion of shRNA seeds for this gene which are associated with this state. E. Predicted targets associated with S5 like states, the labels are similar with D.

For example, for state S4, our approach predicted that knockdown of the proteasome genes *PSMA1* and *PSMB2* would induce a S4-like state (Fig 4D). *PSMB2* is a target of a well-known antineoplastic agent, Carfilzomib. The knockdown of *HDAC3* is also predicted to contribute to the S4-like state, which is consistent with our annotation that S4 is associated with HDAC inhibitors. The method also selected *RUVBL1*, *MCM3*, *MCM7*, *RPS6*, *RPL7* and *CCND1* as being linked to the S4 state. Previous studies have suggested that a selective inhibitor of the RUVBL1/2 complex reduced growth in acute myeloid leukemia and multiple myeloma (Assimon et al., 2019). For state S5, we predicted the knock down of ribosome protein small subunit genes (*RPS15A, RPS16, RPS9, RPS3A, RPS7*) will drive the S5-like state (Fig 4E). The knockdown of *ESR1* also drives the S5-like state, which is consistent with the genetic background of MCF7 as an ER^+^ breast cancer cell line. The method also predicted *SOAT1* as a potential target. *SOAT1* has been investigated in hepatocellular carcinoma, which suggests it is a promising drug target (Jiang et al., 2019). Genes with targetable vulnerability for MCF7-like breast cancer can be found in Fig 4D & 4E and Supplemental Table 2. Our results suggest that the FM-states framework, using integrative analysis of drug induced cell states and functional screening data, can identify potential drug targets.

### Pre-existing transcriptional functional module states are associated with drug response

We next asked whether the transcriptional states of cancer cell lines prior to drug treatment are associated with drug response. We applied the FM-States method to define the cell states using the basal transcriptomic profile from the GDSC project (Iorio et al., 2016), and analyzed the association between cell states and drug response. Using breast cancer cell lines, we generated the FM-factors for all 49 breast cancer cell lines in the GDSC project. We selected potential effective drugs based on absolute IC50 values less than 1 μmol in at least 5 cell lines.

We estimated Spearman correlations between the functional module factors and IC50 values for each selected drug, shown in Fig 5A. The gene expression of cell cycle, replication and repair, metabolism of cofactors and vitamin modules show negative correlations with IC50 for most drugs, suggesting that breast cancer cell lines with higher expression of cell cycle or replication and repair pathway will show higher sensitivity to most drugs if we use a lower IC50 value as the measurement of higher sensitivity (Fig 5B). On the contrary, the gene expression in the transport and catabolism pathway, and carbohydrate metabolism modules show positive correlations with the IC50 of most drugs (Fig 5A & 5C). In contrast, some functional modules such as cell motility, signaling molecules and interaction, signaling transduction modules show different patterns of associations. Cells with overall high expression of these pathways are sensitive to drugs that target the ERK MAPK signaling pathway (anti-correlated), but at the same time are resistant (positive correlated) to drugs that target RTK signaling, PI3K signaling, and chromatin histone acetylation (Fig 5A & 5D). The functional state associated drug response suggests the functional module factors can be used for drug sensitivity prediction.

**Fig 5.**
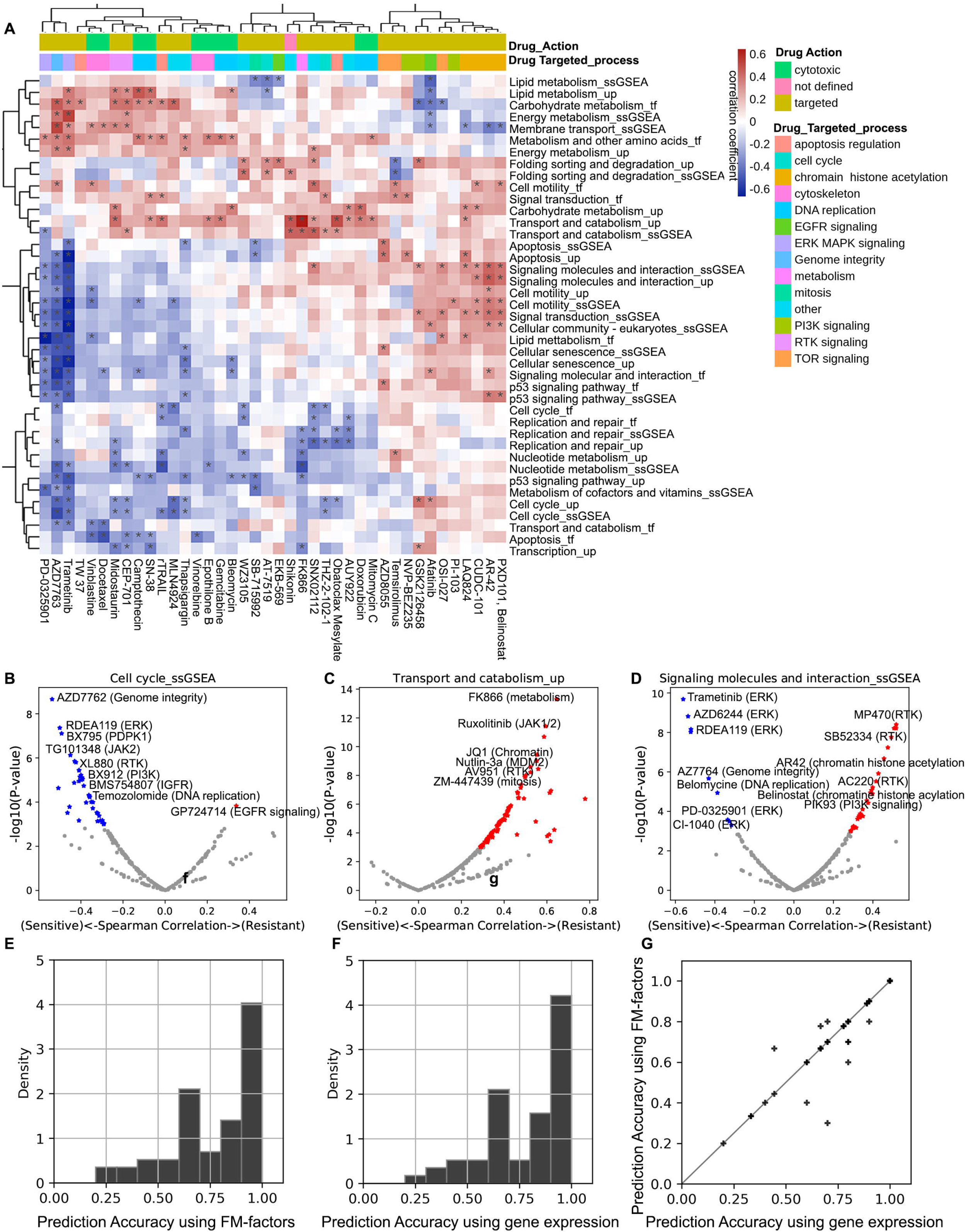
Predicting drug responses using FM-factors. A. Spearman correlation between the FM-factors and drug response (log(IC50)), colored scale by spearman correlation coefficient, * represents significant correlations with threshold *P*-value <0.05. B-D. Volcano plot of Spearman correlation between FM-factors and drug response. The x-axis represents the Spearman correlation coefficient, and the y-axis represents the *P*-values (-log10 transformed). E. Histogram of prediction accuracy of drug response on the testing sets (ratio of samples between training set and testing set is 4:1) using the Random forest model using the FM-factors as features. Each bin displays the bin’s raw count divided by the total number of drugs (n = 57) and the bin width (width = 0.1). F. Distribution of prediction accuracy of drug response using gene expression values as features. G. The comparison of prediction accuracy between the RF models using the FM-factors and all the gene expression values for each drug; each point represents the median prediction accuracy for the 100 bootstrapped RF models using the features of FM-factors and the gene expression values for each drug.

Finally, we wanted to assess whether using functional module factors alone, rather than all gene expression data, would provide comparable performance. We used the Random Forest (RF) algorithm to build predictive models for the sensitivity of cell lines for each drug. The median prediction accuracy for the bootstrapped RF models for all the drugs is 0.8 (Fig 5E). The prediction accuracy using the FM-factor based RF models is comparable with the prediction models using the gene expression data (Fig 5E-G).

## Discussion

The FM-States method considers preliminary knowledge (functional modules or pathways), and defines the FM-factors for any given sample. It helps to generate low dimensional representative features and represents biologically relevant transcriptional states. Further, the FM-States method determines key transcriptional regulators of different states.

Large data sets from consortium projects are rich resources for characterizing the transcriptional states of different cell lines before and after drug treatment or gene knockdown. We showed that FM-States, a simple and biologically interpretable method, is able to predict drug targets and drug response through integrative analysis of multiple data sets. Using the FM-States method, we determined the drug induced transcriptional states for the breast cancer cell line MCF7 using large public gene expression data from the CMap. Different transcriptional states were associated with drug concentration and drug class. We further identified potential targets through integrative analysis of transcriptional states after drug treatment and gene knockdown. By combining the transcriptional states and gene knockdown efficacy, we predicted potential targets for MCF7-like breast cancer, while recognizing the challenge of genetic and non-genetic intra-tumoral heterogeneity. Using the functional module factors as features to represent the transcriptional states, the method revealed transcriptional state dependent drug sensitivity and resistance and exhibited similar performance in a drug response prediction task to a model that uses all gene expression data.

This functional module framework also allows users to extend the analysis to other cell lines or patient derived samples. This work underscores the power of integrating large public data resources to understand and predict drug response.

## Acknowledgements

G. Qin, T. Knijnenburg, and I. Shmulevich are supported by U.S. National Institutes of Health grants U01 CA217883, P01 CA077852 and 5U24CA210952-06. R. Monnat was supported by NCI P01 CA077852. The content is solely the responsibility of the authors, and does not necessarily represent the official views of the National Institutes of Health. We would like to thank the Cancer Target Discovery and Development (CTD^2^) consortium for the discussion and providing resources for this project, the Broad Institute Connectivity Map project (CMap), the Broad Institute Cancer Dependency Map project (DepMap), and the Wellcome Sanger Institute Genomics of Drug Sensitivity in Cancer project (GDSC) for sharing datasets with the research community.

## Author Contributions

Conceptualization and Methodology, G.Qin, T.Knijnenburg and Ilya Shmulevich. Data analysis and implementation: G.Qin. Writing original draft, G.Qin. Writing. Review and editing, I. Shmulevich, D. Gibbs, C. Kemp, R. Moser, R. Monnat, T.Knijnenburg. Supervision, I. Shumulevich.

## STAR Methods

### Selection of functional modules

We consider KEGG processes or pathways as functional modules. The KEGG pathways were annotated in a hierarchical structure, with four main categories: “Metabolism”, “Genetic Information Processing”, “Environmental Information Processing”, and “Cellular Processes”. These categories include the functional gene sets that cover 20 cellular processes: ‘Replication and repair’, ‘Transcription’, ‘Translation’, ‘Folding, sorting and degradation’, ‘Cellular community’, ‘Cell growth and death’, ‘Transport and catabolism’, ‘Cell motility’, ‘Membrane transport’, ‘Signaling transduction’, ‘Signaling molecules and interaction’, ‘Amino acid metabolism’, ‘Metabolism of other amino acids’, ‘Lipid metabolism’, ‘Carbohydrate metabolism’, ‘Metabolism of cofactors and vitamins’, ‘Xenobiotics biodegradation and metabolism’, ‘Glycan biosynthesis and metabolism’, ‘Energy metabolism’, and ‘Nucleotide metabolism’. Each of the 20 cellular processes includes several different pathways, such as the process of ‘cell growth and death’, which includes the pathways ‘Cell cycle’, ‘Apoptosis’, ‘Cellular senescence’, and ‘p53 signaling’. The 23 functional modules in this manuscript include the 4 pathways in the process of cell growth and death and the other 19 cellular processes out of all the 20 processes shown above. These functional modules cover the main functionalities of cells and constitute 5,453 genes and cover most KEGG pathway genes.

### Definition of the FM-factors

To define the FM-factors for each sample, we used four categories of factors: *ssGSEA_score*, *TF_strength*, *up_strength*, and *down_strength*. *ssGSEA_score* is measured using the ssGESA method from the python package of gseapy 0.9.16. The *ssGSEA_score* measures the gene set enrichment score per sample as the normalized difference between the empirical cumulative distribution function of gene expression ranks inside and outside the gene set (Hanzelmann et al., 2013). The *TF_strength* is defined as the average expression of transcription factors whose target genes are enriched in each functional module. It is determined by following these steps:

1. Transcription factor - target gene pairs (TF-Target) that show evidence in either literature-curated resources, ChIP-seq peaks or TF binding motifs on promoters were collected (Garcia-Alonso et al., 2019).
2. TF-target pairs with either curated/high confidence (confidence level A) or likely confidence were selected for further analysis (confidence level B) (Garcia-Alonso et al., 2019).
3. As the states of transcriptional regulatory networks exhibit tissue type or cell type specificity, we further measured the correlation between the transcription factors and their target genes using the input gene expression matrix (e.g., gene expression of MCF7 after drug treatment in the CMap data), and kept only those pairs that showed significant correlation in a given context, such as tumor type (absolute Pearson correlation coefficient >0.2, *P* value <0.05).
4. With the list of TF-target gene pairs, we used the one-tailed Fisher exact test to assess whether the target genes for one TF are enriched in a pathway, and only those TFs that showed significant enrichment in a module were selected as the signature TFs for that module.
5. We then measured the TF regulation *weight* using the following equations (Equations 1–3) for a specific TF, labeled TF_A:

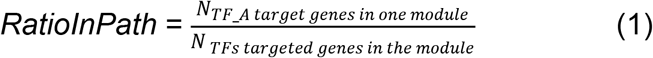

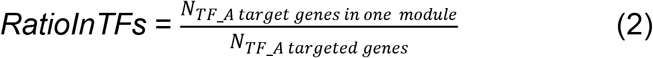

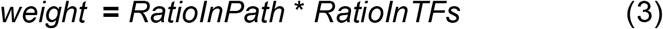

where *N*_*TF _A target genes in a module*_ represents the number of target genes for TF_A in one module, *N*_*TFS target genes in the module*_ represent the number of genes that have been regulated by all the TFs, and *N*_*TF _A target genes*_ represents the number of target genes regulated by TF_A for all selected modules.
6. We normalized the weight in each pathway to make the sum of TF regulation weights equal to 1. We then calculated the average transcriptional strength by summing up the normalized weights, multiplied by the expression level for the master transcription factors (Equation 4), where *M* represents the number of TFs that are estimated to significantly regulate the pathway, and *Expr*(*TF*_*i*_) is the gene expression level for the Transcriptional factor i which regulates genes in the module.

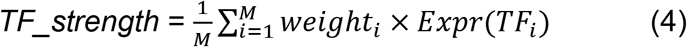

*Up_strength* (*Up-regulation Strength*) and *Down_strength* (*Down-regulation Strength*) are used for estimating the ratio of genes showing high or low expression (|z-score| > threshold) in one functional module for one specific sample compared to all other samples (Equations 5 and 6).

With the absolute gene expression matrix, z-score normalization was performed across all samples for each gene. After the z-score normalization, we define the *Up_strength* and *Down_strength as follows (*Equation 5 and 6):

*Up_strength* for each module is defined as the proportion of up-regulated genes in each module that have z-score above the upper threshold (threshold = 1.6 for our case studies):

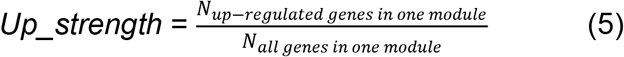

where *N*_*up-regulated genes is on module*_ is the number of up-regulated genes in a module, and *N*_*all genes is on module*_ is the number of genes in that module.

*Down_strength* is defined as the negative fraction of down-regulated genes in a module, with a z-score below the lower threshold:

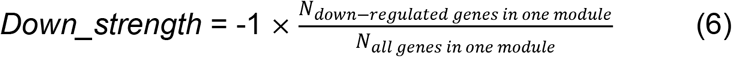

Where *N*_*down-regulated genes is on module*_ is the number of down-regulated genes in a module.

### Annotation of transcriptional states

#### Consensus clustering

We applied the R package of ConsensusClusterPlus (Wilkerson and Hayes, 2010) which uses the consensus clustering method (Monti et al., 2003) to classify the FM-matrix into different clusters, namely FM-states. We select the number of states (clusters) by inspection of the heatmap of the consensus matrix using different ‘*K*’s (K is the number of clusters), the empirical cumulative distribution function (CDF) corresponding to the entries of the consensus matrix, and the relative change in the area under the CDF with the increase of K. When *K*_*true*_ is reached, further increase in the number of clusters does not lead to a corresponding marked increase in the CDF area (Monti et al., 2003).

#### Annotation using the FM-factors

The functional module factors can be easily translated into meaningful biological annotations for each state. To have a better understanding of the functional activity in each state, we detected the key features for each state. Using the Wilcoxon rank-sum test, we tested whether one FM-factor in one state is different from all of the other states. The difference of FM-factors between one state and all the other states are measured using Effect size (Equation 7).

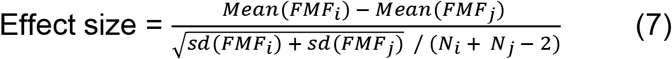

Where *FMF*_*i*_ is the vector of functional module factors for state i, *FMF*_*j*_ is the vector of functional module factors for all the other samples, *N*_*i*_ is the number of samples in states i, and *N*_*j*_ is the number of samples in all the other samples.

We selected the FM-factor for each state showing significant difference (P <0.001 for MCF7 data) and with an effect size greater than a threshold (|effect size| >1 for MCF7 data) between this state and all the other states. The factors ***ssGSEA_score***, ***up_strength***, and ***down_strength*** can be used for the annotation of high or low expression of each functional module for each state. ***TF_strength*** can be used for the annotation of the overall transcriptional regulation strength for each functional module for each state.

#### Annotation using transcriptional factors

To annotate which transcription factors contribute to the regulation of the states, we first identified which transcription factors play activation roles or inhibitory roles for each functional module by 1) selecting the transcription factor -- target gene pairs with positive correlation or negative correlation; 2) selecting transcription factors that show overrepresentation of their target genes in functional modules; 3) identifying transcription factors that show significant (adjusted *P*-value <0.05) differences among different states using the one-way ANOVA test followed by Bonferroni multiple-test correction. Effect size was estimated as the difference of expression of the transcription factor between samples in one state and other states. The generated TF-functional module regulation network was then visualized using Cytoscape (Shannon et al., 2003).

### Defining functional module states for MCF7 cells following drug treatment

FM-factors were generated that cover the four categories of factors for the twenty-three functional modules as described before (see section “Definition of the FM-factors”).

#### 1) Get the relative FM-factors for drug treated samples compared to reference samples

MCF7 transcriptional profiles from the CMap (Subramanian et al., 2017) level 5 data following drug treatment which has reported drug response measurement in the CTRP2 (Rees et al., 2016) were selected, resulting to 1287 drug treated MCF7 profiles. Transcriptional profiles for the DMSO or H20 treated MCF7 samples were selected as reference samples, resulting to 1400 reference samples. Functional module factors (FM-factors) were calculated for both drug treated MCF7 samples and reference MCF7 samples. The FM-factors for drug treated samples are then normalized using the distribution of reference sample FM-factors, which use the ranking method for the *Up-strength*, *Down-strength* and z-score method for *ssGSEA* and *TF_strength*.

#### 2) Clustering of the functional module based factors

Consensus clustering method(Monti et al., 2003; Wilkerson and Hayes, 2010) was used to cluster the FM-factor matrix into stable clusters by inspection of the CDFs’ shape and progression as the number of clusters, K, increases. The most stable classification is considered as the K-solution with the smallest proportion of ambiguous clustering.

#### 3) Annotation

##### 3a) Annotate cell states using the functional module factors

We tested whether a given FM-factor in one state is different from other states using the Wilcoxon rank-sum test. Effect size of difference between one state and all the other states was measured. We selected the FM-factor for each state which showed a significant difference (*P*-value <0.001) and with effect size greater than a threshold (|Effect size| >1) between this state and the other states for the interpretation of each state.

##### 3b) Select the most discriminative transcriptional factors

Activating pairs and inhibitory pairs of transcription factor and target genes were defined by correlation analysis (activating pairs: positive correlation (*P*-value <0.05, cor >0.2), inhibitory pairs: negative correlation (*P*-value <0.05, cor <-0.2)). Transcription factors in the activating TF-target pairs or inhibitory TF-target pairs that show an overrepresentation of their target genes in each functional module were then selected. One way ANOVA test followed by Bonferroni multiple-test correction was then performed to identify whether the transcription factors show a significant (adjusted *P*-value <0.05) difference among different states. We used Cytoscape (Shannon et al., 2003) to visualize the network of transcription factors and functional modules that show differences among different states. The effect size measures the extent to which the TF is high- or low-expressed in each state.

##### 3c) Annotate cell states with external factors

Drug responses (EC50) of MCF7 for different drugs were extracted from the Cancer Therapeutics Response Portal v2 (CTRPv2) (Rees et al., 2016; Seashore-Ludlow et al., 2015). We considered drug concentrations greater than its EC50 values as high concentrations, and drug concentrations smaller than its EC50 values as low concentrations. We measured the ratio of high drug concentration samples to low drug concentration samples and statistically tested whether the distribution of high drug concentration samples and low concentration samples differs among different states using the chi-squared test. The target or action of compounds (drugs) were also from the CTRPv2. Pie plots were used to visualize the distribution of samples with different targets or activity for each state. Fisher’s exact test was used to assess whether one class of drug target is overrepresented in a specific state.

### Use of FM states to predict targetable cancer vulnerabilities

#### Association of gene knockdown to different states

Gene expression profiles of MCF7 after shRNA knockdown are taken from CMap (Subramanian et al., 2017), with each sample treated with one shRNA seed mapping to a specific gene. FM-factors (***ssGSEA_score***, ***TF_strength***, ***up_strength and down_strength***) for all the 23 modules were generated for each sample using the FM-state method. K-nearest neighbors(KNN) classifier (KNeighborsClassifier in the sklearn library in python3) (Pedregosa et al., 2011) was used to assign each sample to one of the defined states after drug treatment. FM-factors (features) which show significant differences (Wilcoxon rank-sum test, *P*-value <0.01 and |effect size| >1) across five states were selected for the KNN models. To select a proper K for the prediction model, we compared the prediction accuracy using five-fold cross validation with K from 1 to 30, increased by 2. K = 5 was selected as the model shows relatively higher prediction accuracy (mean = 0.73) and smaller derivation (standard deviation = 0.05). We then used the KNN model to assign the shRNA treated sample to one of the defined states after drug treatment. Samples predicted as each state with a probability equal to or greater than 0.6 were selected for further analysis and the rest samples with ambiguous prediction results were excluded. As one gene can have different seeds, we further assigned one gene to one state by estimating the enrichment of seeds for this gene in one specific state using Fisher’s exact test. Genes with shRNA seeds enriched (Fisher’s exact test, p <0.05) in one state are assigned to this state.

#### Cancer dependent analysis

Gene knockout effect was downloaded from the DepMap portal (Dempster et al., 2019; DepMap, 2019; Meyers et al., 2017), which contains the results of genome-scale CRISPR knockout screens for 18,333 genes in 625 cell lines. Gene knockout effect data for MCF7 was selected from this dataset. Kolmogorov-Smirnov statistical test was performed to compare the distribution of target genes that drive each state and all genes in the genome-scale CRISPR knockout screen.

#### Selection of potential target genes

For each gene, we calculated the proportion of seeds which are predicted in one state, and we also used the gene knockout effect for the MCF7 cell line. Genes which are predicted to drive a specific state with a gene knockout effect smaller than 0 are selected as target genes that could drive to this state.

### Define basal FM-states of breast cancer cell lines to predict drug sensitivity

We collected gene expression data for all the 49 breast cancer derived cell lines from the GDSC project (Iorio et al., 2016), and applied the FM-States method to this dataset to generate the functional module based factors (FM-factors) for all the 49 breast cancer cell lines. The functional module based factors include *ssGSEA_score*, *TF_strength*, *up_strength, down_strength* for all the 23 pathways were calculated.

#### Correlation analysis between drug response and FM-factors

Drug response data (log(IC50)) was derived from the GDSC datasets (Iorio et al., 2016). As not all of the drugs show efficiency to the breast cancer cell lines, we select potential effective drugs with the filtering criteria: 1) number of cell lines with log(IC50) smaller than zero is greater or equal to 5; 2) number of cell lines tested is greater or equal to 25 (more than half of the samples). Using the selected drugs, we then performed correlation analysis to compare the association between the FM-factors and drug response (log(IC50)) using Spearman correlation.

#### Drug sensitivity prediction models using the FM-factors and gene expression

For each drug, we binarized the drug response for each sample to either sensitive (log(IC50) <-1) or resistant (log(IC50) >-1) response and built a Random Forest (RF) classifier using the python library sklearn (Pedregosa et al., 2011) to predict the drug response using either the feature of FM-factors or gene expression for all the genes. 100 bootstrapped RF models were built, with 80% of the data used for training, 20% for testing. Features importance scores greater than zero are selected for the final models for each drug.

## Data and Code Availability Statements

The code generated during this study is available at https://github.com/IlyaLab/FMStates. All datasets used for the case studies in this study can be found at https://osf.io/34xnm/?view_only=5b968aebebe14d4c97ff9d7ce4cb5070.

## References

Aksoy, B.A., Dancik, V., Smith, K., Mazerik, J.N., Ji, Z., Gross, B., Nikolova, O., Jaber, N., Califano, A., Schreiber, S.L., et al. (2017). CTD2 Dashboard: a searchable web interface to connect validated results from the Cancer Target Discovery and Development Network. Database (Oxford) 2017.

Assimon, V.A., Tang, Y., Vargas, J.D., Lee, G.J., Wu, Z.Y., Lou, K., Yao, B., Menon, M.K., Pios, A., Perez, K.C., et al. (2019). CB-6644 Is a Selective Inhibitor of the RUVBL1/2 Complex with Anticancer Activity. ACS Chem Biol 14, 236–244.

Barretina, J., Caponigro, G., Stransky, N., Venkatesan, K., Margolin, A.A., Kim, S., Wilson, C.J., Lehar, J., Kryukov, G.V., Sonkin, D., et al. (2012). The Cancer Cell Line Encyclopedia enables predictive modelling of anticancer drug sensitivity. Nature 483, 603–607.

Basu, A., Bodycombe, N.E., Cheah, J.H., Price, E.V., Liu, K., Schaefer, G.I., Ebright, R.Y., Stewart, M.L., Ito, D., Wang, S., et al. (2013). An interactive resource to identify cancer genetic and lineage dependencies targeted by small molecules. Cell 154, 1151–1161.

Dempster, J., Rossen, J., Kazachkova, M., Pan, J., Kugener, G., Root, D.E., and Tsherniak, A. (2019). Extracting Biological Insights from the Project Achilles Genome-Scale CRISPR Screens in Cancer Cell Lines. BioRxiv.

DepMap, B. (2019). DepMap 19Q3 Public. figshare. Dataset.

Garcia-Alonso, L., Holland, C.H., Ibrahim, M.M., Turei, D., and Saez-Rodriguez, J. (2019). Benchmark and integration of resources for the estimation of human transcription factor activities. Genome Res 29, 1363–1375.

Ghandi, M., Huang, F.W., Jane-Valbuena, J., Kryukov, G.V., Lo, C.C., McDonald, E.R.3rd,, Barretina, J., Gelfand, E.T., Bielski, C.M., Li, H., et al. (2019). Next-generation characterization of the Cancer Cell Line Encyclopedia. Nature 569, 503–508.

Guo, M., Bao, E.L., Wagner, M., Whitsett, J.A., and Xu, Y. (2017). SLICE: determining cell differentiation and lineage based on single cell entropy. Nucleic Acids Res 45, e54.

Hanzelmann, S., Castelo, R., and Guinney, J. (2013). GSVA: gene set variation analysis for microarray and RNA-seq data. BMC Bioinformatics 14, 7.

Huser, L., Sachindra, S., Granados, K., Federico, A., Larribere, L., Novak, D., Umansky, V., Altevogt, P., and Utikal, J. (2018). SOX2-mediated upregulation of CD24 promotes adaptive resistance toward targeted therapy in melanoma. Int J Cancer 143, 3131–3142.

Iorio, F., Knijnenburg, T.A., Vis, D.J., Bignell, G.R., Menden, M.P., Schubert, M., Aben, N., Goncalves, E., Barthorpe, S., Lightfoot, H., et al. (2016). A Landscape of Pharmacogenomic Interactions in Cancer. Cell 166, 740–754.

Jiang, Y., Sun, A., Zhao, Y., Ying, W., Sun, H., Yang, X., Xing, B., Sun, W., Ren, L., Hu, B., et al. (2019). Proteomics identifies new therapeutic targets of early-stage hepatocellular carcinoma. Nature 567, 257–261.

Jin, S., MacLean, A.L., Peng, T., and Nie, Q. (2018). scEpath: energy landscape-based inference of transition probabilities and cellular trajectories from single-cell transcriptomic data. Bioinformatics 34, 2077–2086.

Kanehisa, M., and Goto, S. (2000). KEGG: kyoto encyclopedia of genes and genomes. Nucleic Acids Res 28, 27–30.

Kim, H.J., and Bae, S.C. (2011). Histone deacetylase inhibitors: molecular mechanisms of action and clinical trials as anti-cancer drugs. Am J Transl Res 3, 166–179.

Kim, J.W., Abudayyeh, O.O., Yeerna, H., Yeang, C.H., Stewart, M., Jenkins, R.W., Kitajima, S., Konieczkowski, D.J., Medetgul-Ernar, K., Cavazos, T., et al. (2017). Decomposing Oncogenic Transcriptional Signatures to Generate Maps of Divergent Cellular States. Cell Syst 5, 105–118 e109.

Knijnenburg, T.A., Wang, L., Zimmermann, M.T., Chambwe, N., Gao, G.F., Cherniack, A.D., Fan, H., Shen, H., Way, G.P., Greene, C.S., et al. (2018). Genomic and Molecular Landscape of DNA Damage Repair Deficiency across The Cancer Genome Atlas. Cell Rep 23, 239–254 e236.

Meyers, R.M., Bryan, J.G., McFarland, J.M., Weir, B.A., Sizemore, A.E., Xu, H., Dharia, N.V., Montgomery, P.G., Cowley, G.S., Pantel, S., et al. (2017). Computational correction of copy number effect improves specificity of CRISPR-Cas9 essentiality screens in cancer cells. Nat Genet 49, 1779–1784.

Monti, S., Tamayo, P., Mesirov, J.P., and Golub, T.R. (2003). Consensus Clustering: A Resampling-Based Method for Class Discovery and Visualization of Gene Expression Microarray Data. Machine Learning 52, 91–118.

Muller, F.U., Loser, K., Kleideiter, U., Neumann, J., von Wallbrunn, C., Dobner, T., Scheld, H.H., Bantel, H., Engels, I.H., Schulze-Osthoff, K., et al. (2004). Transcription factor AP-2alpha triggers apoptosis in cardiac myocytes. Cell Death Differ 11, 485–493.

Ogata, H., Goto, S., Sato, K., Fujibuchi, W., Bono, H., and Kanehisa, M. (1999). KEGG: Kyoto Encyclopedia of Genes and Genomes. Nucleic Acids Res 27, 29–34.

Papaemmanuil, E., Gerstung, M., Bullinger, L., Gaidzik, V.I., Paschka, P., Roberts, N.D., Potter, N.E., Heuser, M., Thol, F., Bolli, N., et al. (2016). Genomic Classification and Prognosis in Acute Myeloid Leukemia. N Engl J Med 374, 2209–2221.

Pedregosa, F., Varoquaux, G., Gramfort, A., Michel, V., Thirion, B., Grisel, O., Blondel, M., Prettenhofer, P., Weiss, R., Dubourg, V., et al. (2011). Scikit-learn: Machine Learning in Python. JMLR 12, 6.

Rees, M.G., Seashore-Ludlow, B., Cheah, J.H., Adams, D.J., Price, E.V., Gill, S., Javaid, S., Coletti, M.E., Jones, V.L., Bodycombe, N.E., et al. (2016). Correlating chemical sensitivity and basal gene expression reveals mechanism of action. Nat Chem Biol 12, 109–116.

Schmidt, M., Fernandez de Mattos, S., van der Horst, A., Klompmaker, R., Kops, G.J., Lam, E.W., Burgering, B.M., and Medema, R.H. (2002). Cell cycle inhibition by FoxO forkhead transcription factors involves downregulation of cyclin D. Mol Cell Biol 22, 7842–7852.

Schmitz, R., Wright, G.W., Huang, D.W., Johnson, C.A., Phelan, J.D., Wang, J.Q., Roulland, S., Kasbekar, M., Young, R.M., Shaffer, A.L., et al. (2018). Genetics and Pathogenesis of Diffuse Large B-Cell Lymphoma. N Engl J Med 378, 1396–1407.

Seashore-Ludlow, B., Rees, M.G., Cheah, J.H., Cokol, M., Price, E.V., Coletti, M.E., Jones, V., Bodycombe, N.E., Soule, C.K., Gould, J., et al. (2015). Harnessing Connectivity in a Large-Scale Small-Molecule Sensitivity Dataset. Cancer Discov 5, 1210–1223.

Shannon, P., Markiel, A., Ozier, O., Baliga, N.S., Wang, J.T., Ramage, D., Amin, N., Schwikowski, B., and Ideker, T. (2003). Cytoscape: a software environment for integrated models of biomolecular interaction networks. Genome Res 13, 2498–2504.

Subik, K., Lee, J.F., Baxter, L., Strzepek, T., Costello, D., Crowley, P., Xing, L., Hung, M.C., Bonfiglio, T., Hicks, D.G., et al. (2010). The Expression Patterns of ER, PR, HER2, CK5/6, EGFR, Ki-67 and AR by Immunohistochemical Analysis in Breast Cancer Cell Lines. Breast Cancer (Auckl) 4, 35–41.

Subramanian, A., Narayan, R., Corsello, S.M., Peck, D.D., Natoli, T.E., Lu, X., Gould, J., Davis, J.F., Tubelli, A.A., Asiedu, J.K., et al. (2017). A Next Generation Connectivity Map: L1000 Platform and the First 1,000,000 Profiles. Cell 171, 1437–1452 e1417.

Teschendorff, A.E., and Enver, T. (2017). Single-cell entropy for accurate estimation of differentiation potency from a cell’s transcriptome. Nat Commun 8, 15599.

Thorsson, V., Gibbs, D.L., Brown, S.D., Wolf, D., Bortone, D.S., Ou Yang, T.H., Porta-Pardo, E., Gao, G.F., Plaisier, C.L., Eddy, J.A., et al. (2018). The Immune Landscape of Cancer. Immunity 48, 812–830 e814.

Trapnell, C., Cacchiarelli, D., Grimsby, J., Pokharel, P., Li, S., Morse, M., Lennon, N.J., Livak, K.J., Mikkelsen, T.S., and Rinn, J.L. (2014). The dynamics and regulators of cell fate decisions are revealed by pseudotemporal ordering of single cells. Nat Biotechnol 32, 381–386.

Troester, M.A., Hoadley, K.A., Parker, J.S., and Perou, C.M. (2004). Prediction of toxicant-specific gene expression signatures after chemotherapeutic treatment of breast cell lines. Environ Health Perspect 112, 1607–1613.

Tsuyuzaki, K., Sato, H., Sato, K., and Nikaido, I. (2020). Benchmarking principal component analysis for large-scale single-cell RNA-sequencing. Genome Biol 21, 9.

Wilkerson, M.D., and Hayes, D.N. (2010). ConsensusClusterPlus: a class discovery tool with confidence assessments and item tracking. Bioinformatics 26, 1572–1573.

Wu, C.H., van Riggelen, J., Yetil, A., Fan, A.C., Bachireddy, P., and Felsher, D.W. (2007). Cellular senescence is an important mechanism of tumor regression upon c-Myc inactivation. Proc Natl Acad Sci U S A 104, 13028–13033.

Zheng, G.X., Terry, J.M., Belgrader, P., Ryvkin, P., Bent, Z.W., Wilson, R., Ziraldo, S.B., Wheeler, T.D., McDermott, G.P., Zhu, J., et al. (2017). Massively parallel digital transcriptional profiling of single cells. Nat Commun 8, 14049.

