## Supplemental Figures and Tables for "A functional module states framework reveals cell states for drug and target prediction"

### Supplemental Information

#### Supplemental Figures

**Fig S1.** Workflow of defining FM-states of MCF7 following drug treatment.

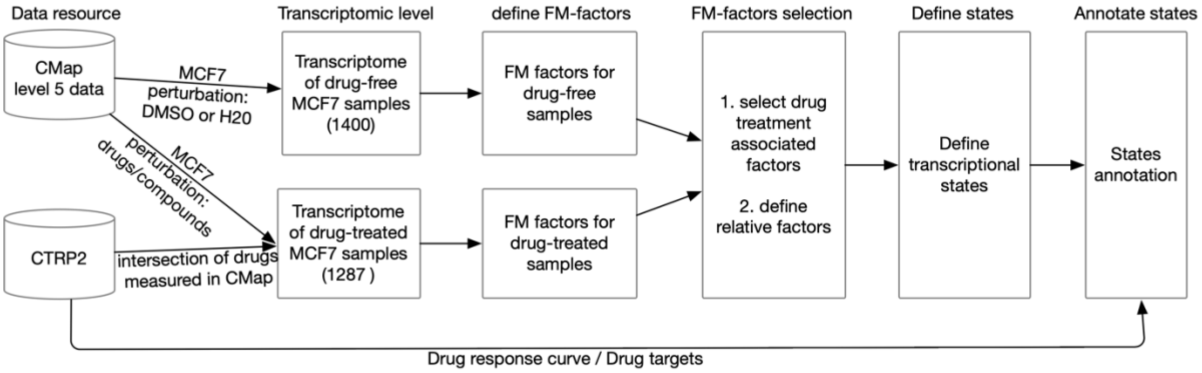

**Fig S2**. Differential FM-factors between drug-treated MCF7 samples and drug-free samples. X-axis represents the effect size of difference of each FM-factor between the drug-treated samples and the drug-free samples, see methods. The Y-axis represents the negative log10 transformed FDR value measured by Mann-Whitney rank test, followed by Benjamini-Hochberg adjustment. Blue color dots show the FM-factors with FDR < 1e-6 and effect size less than -0.2. Red color dots show the FM-factors with FDR < 1e-6 and effect size greater than 0.2.

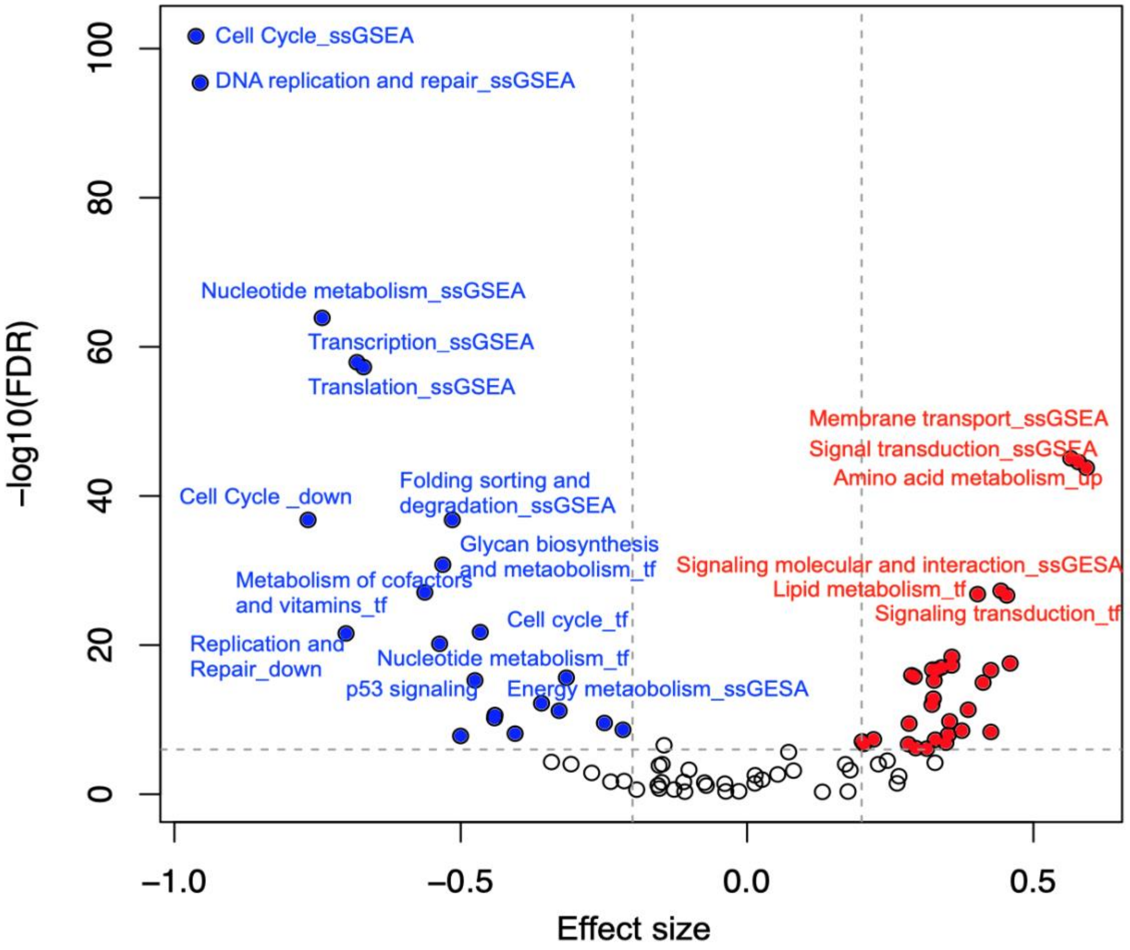

**Fig S3**. Consensus clustering of the relative functional module factors for MCF7 cell lines after drug treatment. A-C. Consensus matrix for the number of clusters k = 3, 4 or 5. D. Cumulative density function for the consensus values. E. Relative change in area under CDF curve with the increase of k. F. Tracking plot of samples with different cluster solutions. Figures are generated using ConsensusClusterPlus R package(Wilkerson and Hayes, 2010).

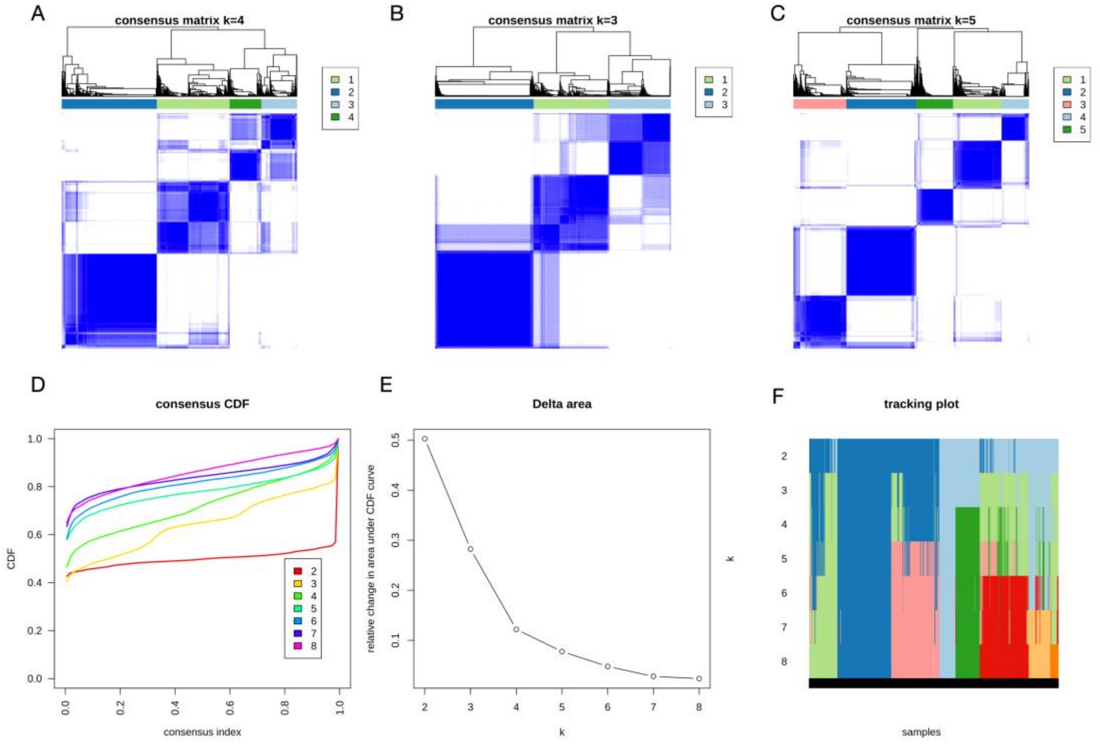

**Fig S4.** The relative expression level of transcriptional factors that show difference among different states.

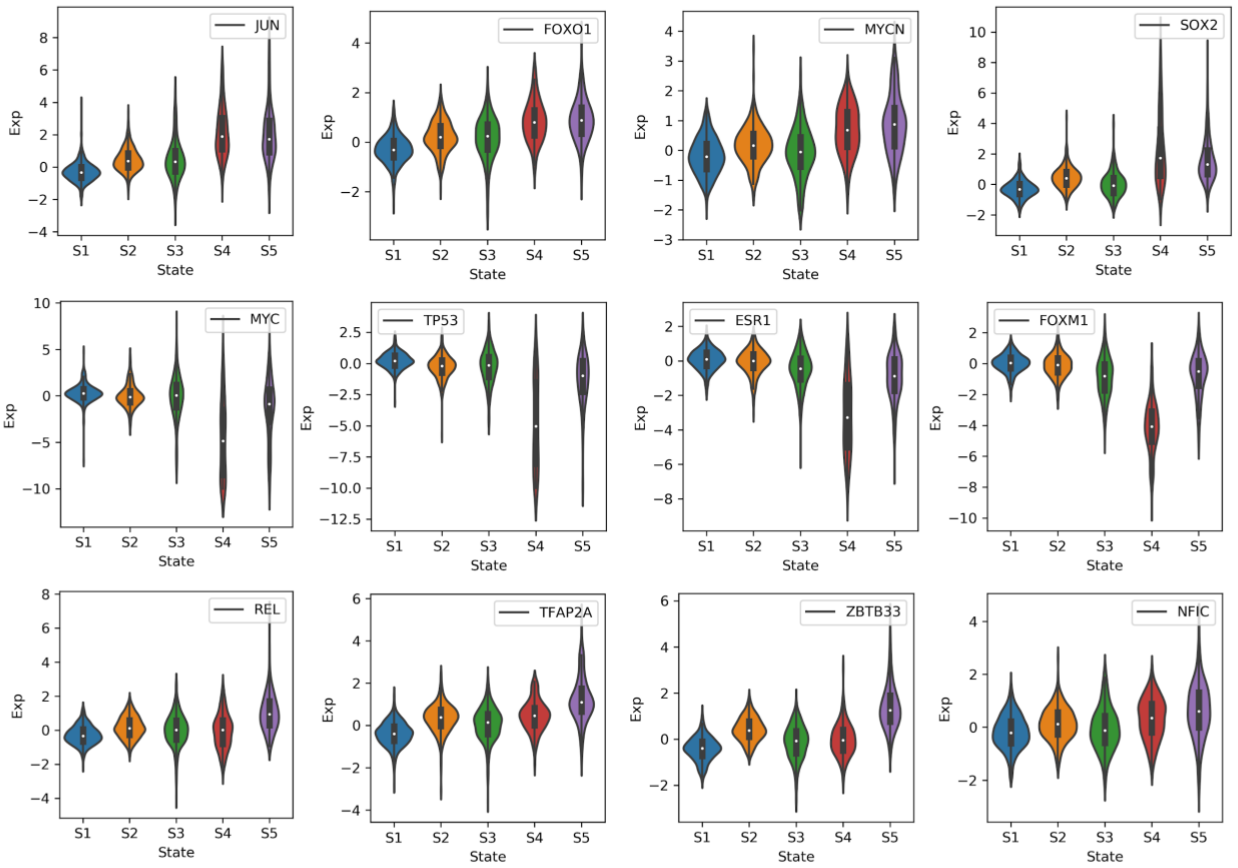

**Fig S5.** Workflow for predicting targetable vulnerabilities using the cell states of MCF7 after drug treatment.

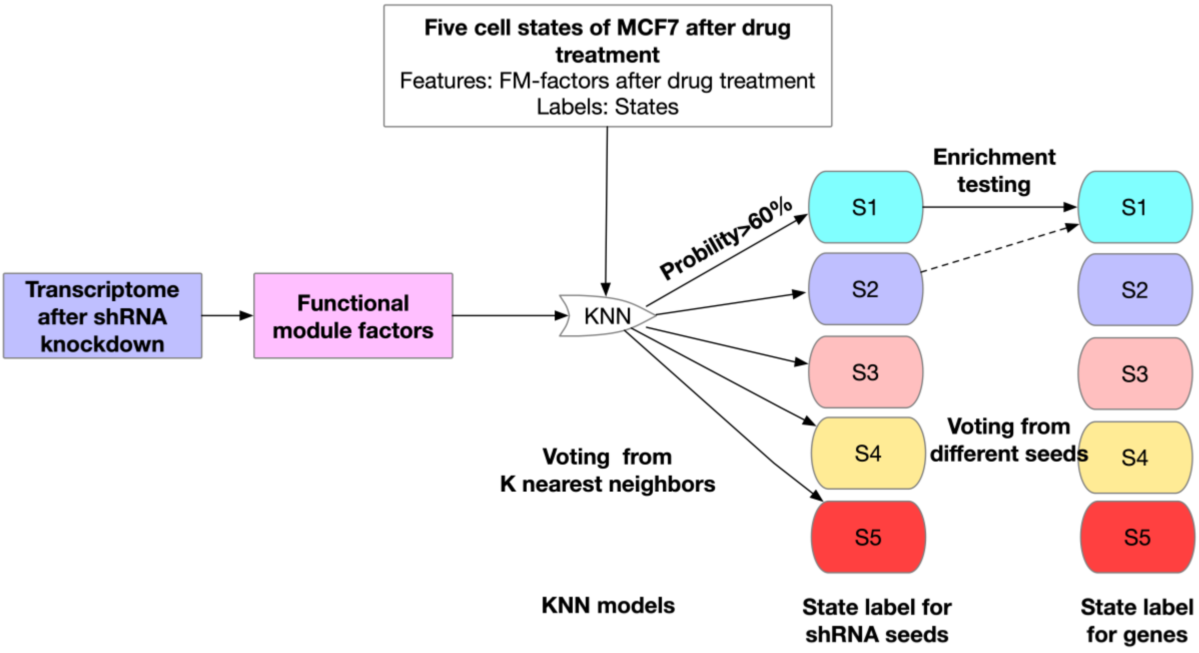

### Supplemental Tables

**Table S1.** Functional modules selected in this study.

| **KEGG Pathways** | ***Category*** |
|---:|---|
| Cellular community - eukaryotes | Cellular Processes |
| Transport and catabolism | Cellular Processes |
| Cell motility | Cellular Processes |
| Apoptosis | Cellular Processes |
| Cellular senescence | Cellular Processes |
| p53 signaling pathway | Cellular Processes |
| Signal transduction | Environmental Information Processing |
| Membrane transport | Environmental Information Processing |
| Signaling molecules and interaction | Environmental Information Processing |
| Translation | Genetic Information Processing |
| Folding sorting and degradation | Genetic Information Processing |
| Replication and repair | Genetic Information Processing |
| Transcription | Genetic Information Processing |
| Cell cycle | Genetic Information Processing |
| Nucleotide metabolism | Metabolism |
| Amino acid metabolism | Metabolism |
| Carbohydrate metabolism | Metabolism |
| Lipid metabolism | Metabolism |
| Metabolism of other amino acids | Metabolism |
| Xenobiotics biodegradation and metabolism | Metabolism |
| Energy metabolism | Metabolism |
| Glycan biosynthesis and metabolism | Metabolism |
| Metabolism of cofactors and vitamins | Metabolism |

**Table S2.** Predicted targetable vulnerability for MCF7-like breast cancer. All genes are predicted to induce the S4 or S5-like state of MCF7, and with gene knockout effects for MCF7 cell line smaller than 0. (Data from Depmap 2019 Q3, Achilles_gene_effect. The lower value of knockout effect represent the higher sensitivity of this gene to the MCF7 cell line). Proportion of seeds are the percentage of shRNA seeds that are predicted to be associated with the states as listed in the right column.

| | Knockout Effect | Proportion of seeds | Associated States |
|---|---|---|---|
| *PSMA1* | -2.26 | 0.50 | S4 |
| *PCNA* | -2.12 | 0.33 | S4 |
| *RUVBL1* | -2.05 | 0.33 | S4 |
| *MCM7* | -1.85 | 0.33 | S4 |
| *CCND1* | -1.85 | 0.20 | S4 |
| *RPL7* | -1.64 | 0.33 | S4 |
| *HSPA9* | -1.42 | 0.33 | S4 |
| *PSMB2* | -1.38 | 0.50 | S4 |
| *HEATR1* | -1.27 | 0.33 | S4 |
| *HDAC3* | -1.22 | 0.33 | S4 |
| *FGFR1OP* | -1.12 | 0.33 | S4 |
| *NDUFS8* | -1.03 | 0.33 | S4 |
| *NFYB* | -0.99 | 0.50 | S4 |
| *PPIE* | -0.89 | 0.33 | S4 |
| *MCM3* | -0.87 | 0.33 | S4 |
| *PFAS* | -0.81 | 0.33 | S4 |
| *ABCF1* | -0.80 | 0.33 | S4 |
| *SMG7* | -0.79 | 0.50 | S4 |
| *COPS2* | -0.79 | 0.33 | S4 |
| *VPS28* | -0.78 | 0.33 | S4 |
| *UBE2Z* | -0.73 | 0.33 | S4 |
| *AGPAT2* | -0.53 | 0.17 | S4 |
| *MOK* | -0.51 | 0.20 | S4 |
| *ZMIZ1* | -0.49 | 0.50 | S4 |
| *PFKL* | -0.47 | 0.33 | S4 |
| *KAT6B* | -0.46 | 0.25 | S4 |
| *MED15* | -0.36 | 0.33 | S4 |
| *UBAP2L* | -0.33 | 0.33 | S4 |
| *CAND1* | -0.33 | 0.33 | S4 |
| *PCSK7* | -0.31 | 0.33 | S4 |
| *FBXO7* | -0.22 | 0.33 | S4 |
| *POLD4* | -0.21 | 0.17 | S4 |
| *FGFR1* | -0.19 | 0.33 | S4 |
| *ZNF785* | -0.19 | 0.33 | S4 |
| *CALM1* | -0.14 | 0.33 | S4 |
| *ZMIZ2* | -0.14 | 0.33 | S4 |
| *SP140* | -0.12 | 0.33 | S4 |
| *TWIST2* | -0.10 | 0.33 | S4 |
| *UGDH* | -0.10 | 0.25 | S4 |
| *ABAT* | -0.09 | 0.20 | S4 |
| *RHOB* | -0.08 | 0.33 | S4 |

| | | | |
|---|---|---|---|
| *PRCP* | -0.06 | 0.50 | S4 |
| *MXD3* | -0.06 | 0.33 | S4 |
| *AKTIP* | -0.06 | 0.33 | S4 |
| *IDH1* | -0.05 | 0.33 | S4 |
| *TSTA3* | -0.04 | 0.33 | S4 |
| *KIF11* | -1.97 | 0.31 | S5 |
| *RPS15A* | -1.83 | 1.00 | S5 |
| *ERH* | -1.71 | 0.67 | S5 |
| *RPS16* | -1.71 | 1.00 | S5 |
| *POLR2A* | -1.68 | 0.67 | S5 |
| *SFPQ* | -1.65 | 0.67 | S5 |
| *RPS9* | -1.49 | 0.67 | S5 |
| *BIRC5* | -1.39 | 0.43 | S5 |
| *SOD1* | -1.28 | 0.55 | S5 |
| *ESR1* | -1.21 | 0.60 | S5 |
| *SENP6* | -1.18 | 0.67 | S5 |
| *SNAPC1* | -1.14 | 0.67 | S5 |
| *ALDOA* | -1.10 | 0.67 | S5 |
| *RPS3A* | -1.01 | 0.67 | S5 |
| *TARDBP* | -0.99 | 0.67 | S5 |
| *RPS7* | -0.95 | 0.67 | S5 |
| *SLC3A2* | -0.93 | 0.67 | S5 |
| *USP7* | -0.88 | 1.00 | S5 |
| *NDUFS3* | -0.84 | 0.67 | S5 |
| *STIL* | -0.81 | 1.00 | S5 |
| *C1QBP* | -0.79 | 0.67 | S5 |
| *SLC35B1* | -0.77 | 0.50 | S5 |
| *QRSL1* | -0.71 | 0.67 | S5 |
| *METAP2* | -0.70 | 0.67 | S5 |
| *HIST1H2BD* | -0.59 | 0.67 | S5 |
| *TERF1* | -0.52 | 0.43 | S5 |
| *ID2* | -0.48 | 0.33 | S5 |
| *SRPRB* | -0.47 | 1.00 | S5 |
| *HMGB1* | -0.46 | 0.67 | S5 |
| *PTK2* | -0.43 | 0.32 | S5 |
| *PET112* | -0.39 | 0.67 | S5 |
| *USP32* | -0.37 | 0.67 | S5 |
| *PSPH* | -0.33 | 0.67 | S5 |
| *NUMB* | -0.31 | 0.67 | S5 |
| *ZFP36L2* | -0.28 | 0.67 | S5 |
| *FGG* | -0.28 | 0.67 | S5 |
| *DCPS* | -0.28 | 0.67 | S5 |
| *KRAS* | -0.27 | 0.19 | S5 |
| *IKBKE* | -0.27 | 0.44 | S5 |
| *ICAM1* | -0.26 | 0.67 | S5 |
| *RXRG* | -0.21 | 0.67 | S5 |
| *AKAP8L* | -0.21 | 1.00 | S5 |
| *SOAT1* | -0.20 | 0.67 | S5 |
| *CSNK1G2* | -0.18 | 0.67 | S5 |
| *RXRA* | -0.18 | 0.67 | S5 |

| | | | |
|---|---|---|---|
| *AKAP11* | -0.18 | 0.67 | S5 |
| *GSDMB* | -0.17 | 0.67 | S5 |
| *MAP2K6* | -0.15 | 0.67 | S5 |
| *PON3* | -0.14 | 0.67 | S5 |
| *IL15* | -0.14 | 0.67 | S5 |
| *PROC* | -0.10 | 0.67 | S5 |
| *PISD* | -0.07 | 0.67 | S5 |
| *RORC* | -0.07 | 0.60 | S5 |
| *CNDP2* | -0.07 | 0.67 | S5 |
| *NUCB2* | -0.05 | 0.67 | S5 |
| *B4GALT4* | -0.03 | 0.67 | S5 |
| *ZNF662* | -0.03 | 0.67 | S5 |
| *AKR1A1* | -0.03 | 0.75 | S5 |
| *NEK7* | -0.02 | 0.67 | S5 |
| *LGR6* | -0.02 | 0.67 | S5 |
| *EIF2AK3* | -0.01 | 0.40 | S5 |
| *BPHL* | -0.01 | 0.67 | S5 |

**References**:

Wilkerson, M.D., and Hayes, D.N. (2010). ConsensusClusterPlus: a class discovery tool with confidence assessments and item tracking. Bioinformatics *26*, 1572-1573.
